# Optimized transcriptional signature for evaluation of MEK/ERK pathway baseline activity and long-term modulations in ovarian cancer

**DOI:** 10.1101/2022.03.21.485160

**Authors:** Mikhail S. Chesnokov, Anil Yadav, Ilana Chefetz

**Affiliations:** The Hormel Institute, University of Minnesota, Austin, Minnesota, USA; Masonic Cancer Center, Minneapolis, Minnesota, USA; Stem Cell Institute, Minneapolis, Minnesota, USA

## Abstract

Ovarian cancer is the most aggressive and lethal of all gynecologic malignancies. High activity of the MEK/ERK signaling pathway is tightly associated with tumor growth, high recurrence rate, and treatment resistance. Several transcriptional signatures were proposed recently for evaluation of MEK/ERK activity in tumor tissue. In the present study, we validated the performance of a robust multi-cancer MPAS 10-gene signature in various experimental models and publicly available sets of ovarian cancer samples. Expression of four MPAS genes (*PHLDA1, DUSP4, EPHA2*, and *SPRY4*) displayed reproducible responses to MEK/ERK activity modulations across several experimental models *in vitro* and *in vivo*. Levels of *PHLDA1, DUSP4*, and *EPHA2* expression were also significantly associated with baseline levels of MEK/ERK pathway activity in multiple human ovarian cancer cell lines and ovarian cancer patient samples available from the TCGA database. High *EPHA2* expression, platinum therapy resistance, and advanced age at diagnosis were associated with poor overall patient survival. Taken together, our results demonstrate that performance of transcriptional signatures is significantly affected by tissue specificity and aspects of particular experimental models. We therefore propose that gene expression signatures derived from comprehensive multi-cancer studies should be always validated for each cancer type.

## INTRODUCTION

Ovarian cancer is the 5^th^ leading cause of death among all cancers in women, with a mortality rate exceeding 60% (1). Most ovarian cancers are high-grade serous ovarian carcinomas (HGSOC), an epithelial tumor subtype that is distinguished by its high aggressiveness, lack of approaches for early diagnosis, and therefore poor prognosis (2–4). Current HGSOC treatment strategies revolve around administration of platinum-based chemotherapeutic agents (cisplatin, carboplatin, etc.) that initially are effective in most cases. Subsequent relapse and acquired treatment resistance remain a major challenge (5–8). Other drugs affecting cancer cells in either a systemic (taxanes, doxorubicin) or targeted way (inhibitors of poly-(ADP-ribose)-polymerase, vascular endothelial growth factor, topoisomerase) are used to bypass platinum resistance, but eventually ovarian tumors develop multi-drug resistance through various molecular mechanisms (9). It is therefore very important to establish new treatment approaches that would target either known resistance-associated genes or new pro-oncogenic regulatory pathways.

While many regulatory circuits involved in ovarian cancer drug resistance were known for a long time (9), new potentially targetable pathways are being revealed based on genomic, epigenomic, and transcriptomic data analysis (10). One of them is the mitogen-activated protein kinase 1/2 (MEK1/2)-extracellular signal-regulated kinase 1/2 (ERK1/2) branch of the mitogen-activated protein kinase (MAPK) pathway, a crucial proliferation and survival regulator in most cells, especially malignant ones (11, 12). While the MEK/ERK pathway in tumors has been thoroughly studied (12, 13), its role in ovarian cancer is just now being assessed. Hyperactivation of MEK1/2 and ERK1/2 is often caused by activating mutations in Kirsten rat sarcoma viral oncogene homolog (*KRAS*) and V-Raf murine sarcoma viral oncogene homolog B (*BRAF*) genes, and therefore was considered to be confined to low-grade ovarian tumors only (14, 15). However, recent analyses of accumulated genomic data revealed that HGSOC, while rarely harboring *KRAS*/*BRAF* mutations, still often displays amplifications and overexpression of MEK1/2 pathway elements (16). High MEK1/2 activity was experimentally confirmed in HGSOC cell lines and clinical samples and identified as a negative prognostic factor (17, 18). MEK1/2 activation occurs in response to cisplatin and may render tumors resistant to platinum treatment (18–20). Moreover, the MEK1/2 pathway affects the subpopulation of chemoresistant ovarian cancer stem-like cells (CSCs) defined by high ALDH1 levels (18, 21–23). Taken together, these data clearly support the importance of the MEK1/2-ERK1/2 pathway in HGSOC and its high potential as a therapeutic target, since it is both a regulator of HGSOC development and a pathway involved in drug resistance.

Direct detection of total or active (e.g. phosphorylated) protein is a reliable way to evaluate signaling pathway activity, but is challenging to perform in small samples (for example, obtained via fine needle aspiration). Moreover, the detectable level of regulatory protein itself, its real activity level, and its effects on downstream targets are not always tightly connected (24, 25). For the MEK1/2 pathway, response to MEK inhibitors cannot be reliably predicted based on evaluation of phosphorylated ERK1/2 alone (26). It is therefore useful to define specific signatures of genes/proteins regulated by pathways of interest, as more robust and reliable indicators of signaling pathway activity than single-molecule biomarkers.

An overwhelming number of genes associated with KRAS/BRAF/MEK/ERK activity have been reported in the literature (summarized in (26)). Attempts to discern a simpler and more relevant MEK1/2-specific signature have been made recently. Two 18-gene and 13-gene signatures predicted sensitivity to the selective MEK1/2 inhibitor selumetinib in a multi-type cancer cell line panel (26) and were independently confirmed in a set of gastric cancer cell lines (27). A set of 24 genes encoding MEK1/2 pathway proteins displayed significant prognostic potential and associations with metabolic aberrations in hepatocellular carcinoma (28). In addition, a 20-gene signature that predicted survival was identified in cisplatin-sensitive HGSOC based on proteomic and transcriptomic data (17).

Recently, a robust 10-gene signature, the MAPK Pathway Activity Score (MPAS), was described as a potent tool to predict the sensitivity of multiple different tumor cells to MEK inhibition (29). Instead of being identified through analysis of large transcriptomic datasets, this MEK-specific signature was built based on manual selection of direct MEK/ERK targets reported in multiple types of cancer. High MPAS was associated with higher sensitivity to MEK1/2 inhibition by cobimetinib in multiple malignant cell lines, increased response to the BRAF inhibitor vemurafenib in BRAF^V600^-positive melanoma patients, and negative survival prognosis. Earlier we successfully utilized MPAS genes to confirm MEK1/2-ERK1/2 pathway inhibition by trametinib in HGSOC cell lines and xenografts (18). In the present study, we aimed to further refine the MPAS gene signature and improve its efficiency in HGSOC by functionally linking MEK1/2-ERK1/2 activity to background levels of MPAS genes and changes, using HGSOC cell lines and patient samples from The Cancer Genome Atlas database.

## MATERIALS AND METHODS

### Cell cultures

We used 20 different human ovarian cancer cell lines, including Type I and Type II ovarian cancer cells (see Table S1 for details). OVCAR8, PEO4, and A2780 cells were provided by Dr. S. Murphy (Duke University, Durham, NC, USA). OVCAR3, OVCAR5, OVCAR7, OVCAR10, OVSAHO and ES2 cell lines were provided by Dr. V. Shridhar (Mayo Clinic, Rochester, MN, USA). Pt152 and Pt486 cells were provided by Dr. R. J. Buckanovich (University of Pittsburgh, Pittsburgh, PA, USA). The Kuramochi cell line was purchased from the Japanese Collection of Research Bioresources Cell Bank (Japan). CAOV3, SKOV3, NIH-OVCAR3, TOV112D (also known as TOV21D), HEY, and TOV21G cells were purchased from American Type Culture Collection (ATCC, Manassas, VA, USA). COV362 and PEO1 cells were purchased from European Collection of Authenticated Cell Cultures (Millipore Sigma, Burlington, MA, USA).

All cell lines were cultivated in RPMI-1640 medium (Corning, Tewksbury, MA, USA) supplemented with 5% FBS (Thermo Fisher Scientific, Waltham, MA, USA) and 100 U/mL penicillin/100 μg/mL streptomycin (Corning, Tewksbury, MA, USA). PEO1 cells were cultivated in medium with the addition of 1 mM sodium pyruvate (Corning, Tewksbury, MA, USA). Cells were tested for mycoplasma monthly.

### Drug treatment of cultured cells

FGF4 (STEMCELL Technologies, Vancouver, Canada) was dissolved in sterile water. Aphidicolin (Cayman Chemical, Ann Arbor, MI, USA) and trametinib (Selleck Chemicals, Houston, TX, USA) were dissolved in DMSO (Fisher Scientific, Waltham, MA, USA). Cells were seeded to the culture dishes and allowed to adhere to the surface overnight. Cell growth media were aspirated and replaced with treatment media containing the drug of interest. Control samples in all experiments performed were treated with vehicle only. Vehicle concentration in growth medium did not exceed 0.2%.

### Synchronization of cultured cells in different phases of cell cycle

Cells were cultured in FBS-free growth medium for 24 hours and subsequently treated with 2 μg/mL aphidicolin in full growth medium for 20 hours to induce cell cycle arrest at early S-phase. Aphidicolin-treated cells were washed twice with sterile DPBS (Corning, Tewksbury, MA, USA) and either harvested immediately or cultivated in full growth medium for 1.5, 3, 6, 10, or 27 hours before harvesting.

### Cell cycle assays

Cells were harvested by trypsinization, washed once with ice-cold full growth medium, and centrifuged at 200 g for 5 minutes. Cells were resuspended in 300 μL of ice-cold PBS and fixed by the addition of 700 μL of ice-cold 70% ethanol in a dropwise manner with constant mild vortexing. Fixed samples were incubated at −20°C overnight. Fixed cell samples were washed with ice-cold PBS (Genesee Scientific, El Cajon, CA, USA) twice, treated with 0.2 mg/mL RNAse A (Thermo Fisher Scientific, Waltham, MA, USA) in PBS for 60 minutes at 37°C, and stained with 10 µg/mL propidium iodide (Millipore Sigma, Burlington, MA, USA) in PBS. Stained samples were analyzed using an FACSCalibur flow cytometer (BD Biosciences, San Jose, CA, USA) and ModFit LT software (Verity Software House, Topsham, ME, USA).

### RT-qPCR analysis of gene expression

Cells were harvested by trypsinization, washed with ice-cold PBS, and centrifuged at 200 g for 5 minutes. Total RNA was isolated from cell pellets using TRIzol reagent and PureLink RNA Mini Kit (Thermo Fisher Scientific, Waltham, MA, USA) with on-column DNAse treatment. A RevertAid RT Reverse Transcription Kit (Thermo Fisher Scientific, Waltham, MA, USA) with a combination of oligo(dT)_18_ and random primers was used to generate cDNA from 1 μg of total RNA. Quantitative PCR analysis of gene expression was performed in a CFX96 Touch thermocycler (Bio-Rad Laboratories, Hercules, CA, USA) using PowerUp SYBR Green Master Mix (Thermo Fisher Scientific, Waltham, MA, USA) and primers listed in Table S2. A three-step amplification program (15 seconds at 95°C, 45 seconds at 62°C, 30 seconds at 72°C) was run for 40 cycles and reaction specificity was checked by melt curve analysis and agarose gel electrophoresis. Reaction efficiency was evaluated using a standard curve approach and was within 98-102% for all primers. Transcript abundance was estimated using Pfaffl’s approach (30). *TBP* was used as a housekeeping normalization gene.

### Immunoblotting

Cells were harvested by scraping in ice-cold PBS and centrifuged at 200 g for 5 minutes. Total protein extracts were obtained from cell pellets using Pierce RIPA buffer (Thermo Fisher Scientific, Waltham, MA, USA) supplemented with Halt Protease and Phosphatase Inhibitor Cocktail (Thermo Fisher Scientific). Protein concentrations were estimated using a Pierce BCA Protein Assay Kit (Thermo Fisher Scientific). 40 μg of total protein was separated in Bolt 4-12% Bis-Tris Plus Gels (Thermo Fisher Scientific) and transferred to Hybond P 0.45 PVDF membranes (GE Healthcare, Chicago, IL, USA). Membranes were blocked with 5% BSA (Thermo Fisher Scientific) in tris-buffered saline with 0.1% Tween-20 (TBST) (Fisher Scientific) and probed overnight at 4°C with the following primary antibodies diluted in 5% BSA in TBST: pMEK1/2 (Cell Signaling Technology, Danvers, MA, USA, 41G9, 1:1000), pERK1/2 (Cell Signaling Technology, D13.14.4E, 1:1000), total ERK1/2 (Cell Signaling Technology, 137F5, 1:1000), pp90RSK1 (R&D Systems, Minneapolis, MN, USA, 1024A, 1:1000), GAPDH (ProteinTech, Rosemont, IL, USA, 1E6D9, 1:10000). Secondary antibodies were HRP-conjugated anti-rabbit IgG (Jackson Immunoresearch, West Grove, PA, USA, 1:10000) or anti-mouse IgG (Jackson Immunoresearch, 1:10000) diluted in 5% skim milk (Millipore Sigma, Burlington, MA, USA) in TBST. Protein bands were developed using Luminata Classico or Luminata Forte HRP substrate (Millipore Sigma) and detected using an Amersham Imager 600 (GE Healthcare, Chicago, IL, USA). After band detection, every membrane was incubated in Restore PLUS Western Blot Stripping Buffer (Thermo Fisher Scientific) for 30 minutes at room temperature and re-probed. Membranes previously stained for pERK1/2 were re-probed for tERK1/2, and other membranes were re-probed for GAPDH. Densitometric analysis of immunoblot images was performed using Image Lab V 6.1.0 software (Bio-Rad Laboratories, Hercules, CA, USA). Raw band intensity values obtained for proteins of interest were divided to corresponding intensity values of GAPDH bands detected using the same immunoblot membrane. Resulting values were additionally normalized to one of the samples, with relative band intensity level for this sample being equal to 1.00 (see Figure Legends for detailed description of each immunoblot experiment).

### Cell viability assays

Cells were plated in 12-well plates (Olympus Plastics, El Cajon, CA, USA) at 50,000 cells/well and synchronized in different phases of the cell cycle as described above. Cells enriched in the cell cycle phase of interest were treated with 100 nM trametinib for 72 hours. Treated cells were harvested by trypsinization, pelleted, and resuspended in PBS. Numbers of viable and dead cells were assessed by direct counting using a Countess II automated cell counter (Thermo Fisher Scientific) in the presence of 0.4% Trypan Blue.

### Analysis of TCGA datasets

Two publicly available datasets from the TCGA project (https://www.cancer.gov/tcga) were used: the “Nature 2011” dataset (31) and the “PanCancer Atlas” dataset (32). Protein RPPA data (median-centered normalized values) were obtained from the “PanCancer Atlas” dataset. Transcriptomic microarray data (Z-scores) were obtained from the “Nature 2011” dataset. Clinical data were combined from information available in both the “PanCancer Atlas” and “Nature 2011” datasets.

### Statistics

Statistical tests used for evaluation of difference between experimental groups are mentioned in the corresponding Figure Legends. Z-score values were calculated by subtracting sample mean values from the individual sample values and then dividing the result to the sample standard deviation. Unsupervised hierarchical cluster analysis was performed using Morpheus software (https://software.broadinstitute.org/morpheus) with the “furthest neighbor” algorithm and either 1-minus Spearman’s rank correlation or Euclidean distance metrics (mentioned in the corresponding Figure Legends). Correlation analysis was performed in OriginPro 2016 software (OriginLab Corporation, Northampton, MA, USA) using a Spearman’s rank correlation approach. Survival analysis was performed in OriginPro 2016 software (OriginLab Corporation) using either the Kaplan-Meyer estimator model with log-rank test or the Cox proportional hazards model. Graphs were plotted using Prism 8 software (GraphPad Software, San Diego, CA, USA) and OriginPro 2016 software (OriginLab Corporation).

## RESULTS

### PHLDA1, SPRY4, EPHA2, and DUSP4 were the best MEK/ERK responders among ten MPAS genes

Expression of the MPAS gene signature is associated with MEK/ERK pathway activity across various tumors (29). Therefore, we used it to confirm MEK/ERK inhibition by trametinib in ovarian cancer (18). Most MPAS genes displayed prominent downregulation in response to trametinib, but two genes (*DUSP6* and *CCND1*) displayed inconsistent results, and *EPHA4* expression levels were below reliable levels of detection. The original MPAS study demonstrated that contribution of single genes to the final MPAS signature can be tissue-specific, so we optimized this approach for ovarian cancer samples.

From the functional point of view, the best transcriptional indicators of signaling pathway activity should be among the genes predominantly controlled by the pathway of interest. We therefore analyzed the response of MPAS genes to MEK/ERK activity modulation in PEO4 cells *in vitro* (Figure 1). We chose PEO4 cells as a primary experimental model because they display moderate MEK/ERK activity at baseline, and we obtained data about some of their MEK/ERK-related aspects in our earlier report (18).

**Figure 1.**
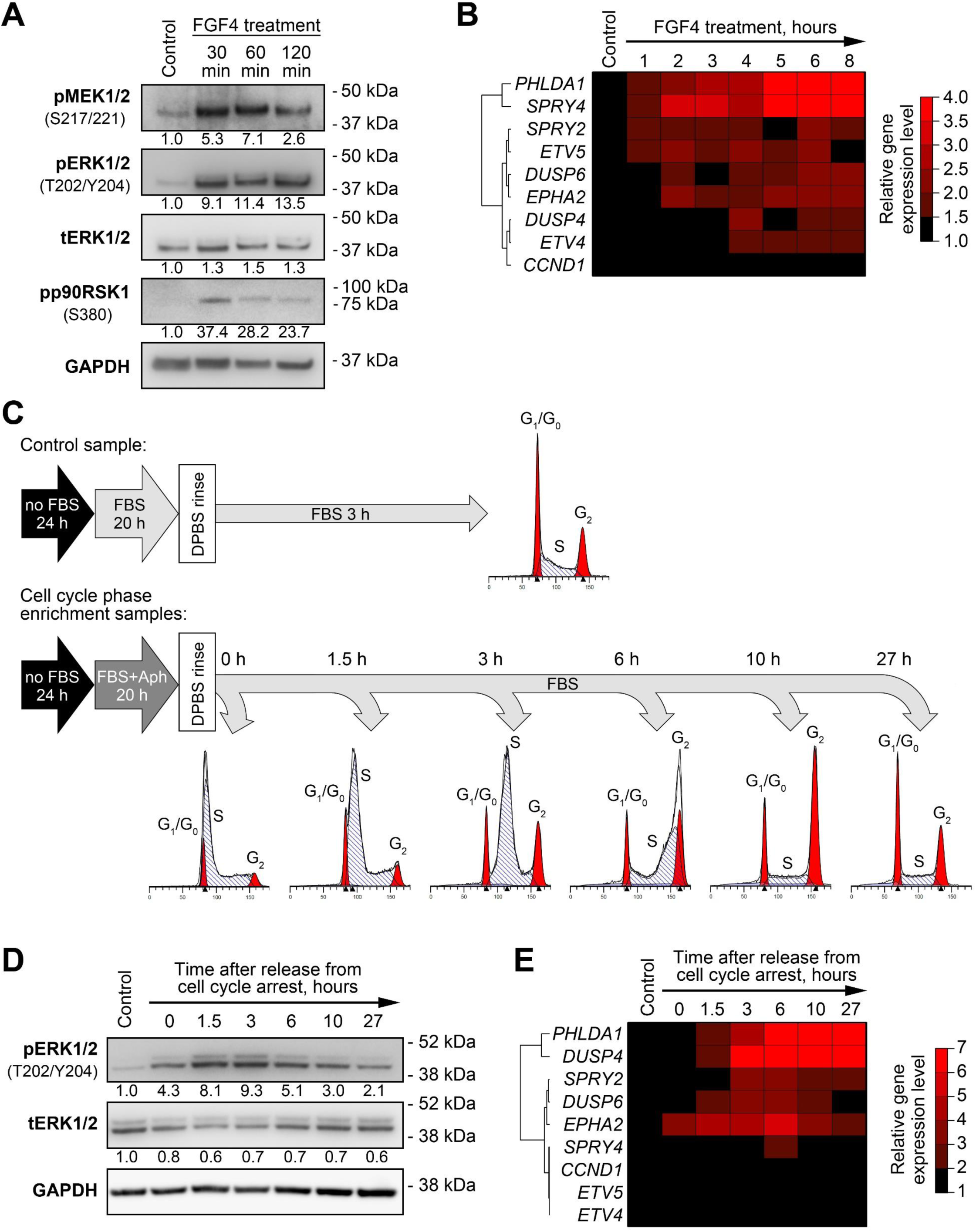
Changes in expression levels of MPAS genes induced by MEK/ERK pathway activity modulation in PEO4 cells. (A) Immunoblotting analysis of phosphorylation of MEK/ERK pathway components in response to treatment with FGF4 (100 ng/mL). Numbers under the bands represent relative intensity normalized to GAPDH levels and control samples. (B) Changes in expression of MPAS genes in FGF4-treated cells. Heatmap represents gene expression levels normalized to control samples. Unsupervised hierarchical clustering was performed using 1-minus Spearman’s rank correlation metrics. (C) Experimental design for enrichment of cultured cells in different phases of cell cycle (see Methods for details). Aph – aphidicolin. (D) Immunoblotting analysis of ERK1/2 phosphorylation changes during cell cycle progression in synchronized cells. Numbers under the bands represent relative intensity normalized to GAPDH levels and control samples. (E) Changes in expression of MPAS genes during cell cycle progression. Heatmap represents gene expression levels normalized to control samples. Unsupervised hierarchical clustering was performed using 1-minus Spearman’s rank correlation metrics.

Fibroblast growth factor 4 (FGF4) binds to its receptors on the cell membrane and stimulates activation of various signaling pathways, including the MEK/ERK pathway (33). As expected, treatment of cells with FGF4 resulted in prominent increases in phosphorylation of MEK1/2, ERK1/2, and their downstream target P90RSK within 30 minutes after treatment started (Figure 1A). Changes in expression patterns of MPAS genes in FGF4-treated cells were diverse. *PHLDA1* and *SPRY4* were strongly upregulated (up to a 5-to 6-fold increase after 6 hours); *DUSP6, EPHA2*, and *ETV5* levels increased 2-fold; and *SPRY2, DUSP4, ETV4*, and *CCND1* displayed no substantial changes (Figure 1B, Figure S1). As in our previous report (18), *EPHA4* levels in both control and treated PEO4 cells were below the limit of reliable detection with RT-qPCR, so we excluded this gene from further analysis. *PHLDA1, SPRY4, EPHA2*, and *ETV5* genes responded to MEK/ERK activation relatively quickly, as their levels were upregulated 2-fold or higher within 2 hours after treatment began, while *DUSP6* reached a threshold (2-fold expression increase) after 6 hours (Figure 1B, Figure S1). The changes described above are summarized in Table S3.

MEK/ERK pathway activity in tumors, including HGSOC, is tightly connected to cell proliferation and is required for G1-to-S-phase transition in the cell cycle (18). We therefore hypothesized that changes in MEK/ERK activity and MPAS gene expression during cell cycle progression may be used as a more physiologically relevant model. We used aphidicolin treatment with subsequent release to synchronize PEO4 cells and enrich them in different phases of the cell cycle (Figure 1C). ERK1/2 activity was strongly elevated during early and mid-S phase (0-3 hours timepoints) and then slowly returned to original levels while the cells progressed through G2 and M phases (Figure 1D). Similar to the FGF4 experiment, we could separate MPAS genes into three groups based on their response to ERK1/2 changes: strong response (*PHLDA1, DUSP4*), moderate response (*EPHA2, SPRY2, DUSP6*), and weak/no response (*SPRY4, ETV4, ETV5, CCND1*) (Figure 1E, Figure S2). *PHLDA1, DUSP4, EPHA2*, and *SPRY2* genes exhibited lasting changes, still displaying more than 2-fold increases in their expression levels 27 hours past the release from S phase arrest (Figure 1E, Figure S2; see Table S3 for the summary).

To select the best MEK/ERK responder genes, we assigned points to each evaluated gene based on its performance in the experiments described above and in our previous report (18). The final score obtained allowed us to identify the most reliable responders to MEK/ERK signaling changes (Table 1). Most genes we examined responded robustly to MEK inhibition both *in vitro* and *in vivo*, but only some of them displayed reproducible upregulation upon MEK/ERK pathway activation across different models. We therefore concluded that the whole MPAS gene signature exhibited clear redundancy in HGSOC and focused our further analysis on four genes displaying the most robust response to MEK/ERK activity modulation – *PHLDA1, SPRY4, EPHA2*, and *DUSP4*, henceforth referred to as “Carcinoma of the Ovary MEK/ERK Signature” (COMS).

**Table 1.** Summary of MPAS genes’ response to MEK-ERK pathway activity changes across all analyzed experimental models (including data reported in Chesnokov et al., 2021).

| Gene | Response to MEK-ERK activity changes |  |  |  |  | Total score |
| --- | --- | --- | --- | --- | --- | --- |
|  | Inhibition by trametinib (Chesnokov et al., 2021)<sup>a</sup> |  | Activation by FGF4<sup>b</sup> | Modulation during cell cycle progression<sup>c</sup> | Activation in cisplatin-resistant cells (Chesnokov et al., 2021)<sup>d</sup> |  |
|  | In vitro | In vivo |  |  |  |  |
| *PHLDA1* | 2 | 2 | 2 | 2 | 2 | 10 |
| *SPRY4* | 2 | 2 | 2 | 1 | 0 | 7 |
| *DUSP4* | 2 | 1 | 0 | 2 | 2 | 7 |
| *EPHA2* | 2 | 1 | 1 | 1 | 1 | 6 |
| *DUSP6* | 0 | 2 | 1 | 1 | 1 | 5 |
| *SPRY2* | 2 | 0 | 0 | 1 | 2 | 5 |
| *ETV4* | 2 | 1 | 0 | 0 | 0 | 4 |
| *ETV5* | 2 | 1 | 1 | 0 | 0 | 4 |
| *CCND1* | 2 | 0 | 0 | 0 | 0 | 2 |

<sup>a</sup>Scores are calculated as follows:
- More than 2-fold decrease – 2 points; - 1.5-2-fold decrease – 1 point;
<sup>b</sup>Scores are calculated as follows:
- Strong response – 2 points; - Moderate response – 1 point;
<sup>c</sup>Scores are calculated as follows:
<sup>d</sup>Scores are calculated as follows:
- More than 4-fold increase – 2 points; - 2-4-fold increase – 1 point;

### PHLDA1, EPHA2, and DUSP4 are associated with baseline MEK/ERK activity across multiple ovarian cancer cell lines

Prominent gene expression response to various external or internal stimuli is useful for tracking pathway modulation in experimental models. On the other hand, gene expression patterns in clinical samples usually represent either a baseline state or long-term processes. Thus, gene signatures that reflect baseline levels of pathway activity have much higher potential for clinical applicability.

We evaluated possible associations between expression of COMS genes and MEK/ERK pathway activity in 20 ovarian cancer cell lines. Phosphorylation levels of MEK1/2, ERK1/2, and P90RSK in these cells were evaluated using immunoblotting. Five cell lines were excluded from further analysis based on discrepancies between individual elements of MEK/ERK pathway (e.g. high pERK1/2 without pMEK1/2 or high pP90RSK with no pERK1/2) (Figure 2A, Figure S3). Individual cell lines displayed marked differences in both protein-estimated MEK/ERK activity and expression levels of COMS genes (Figure 2, Figure S4). We therefore analyzed possible correlations between expression levels of COMS genes and relative levels of pERK1/2 normalized to total ERK1/2 expression, using unsupervised hierarchical clustering and Spearman’s correlation approaches. Three genes (*PHLDA1, DUSP4*, and *EPHA2*) displayed positive associations with ERK1/2 phosphorylation levels in both tests, while *SPRY4* exhibited no statistically significant association (Figure 2B, Table 2).

**Table 2.** Associations between expression levels of the most promising MEK/ERK responders and baseline ERK phosphorylation levels in ovarian cancer cell lines.

| Expression levels of MEK/ERK-responding genes |  | Relative ERK1/2 phosphorylation level |
| --- | --- | --- |
| *PHLDA1* | Spearman's correlation coefficient | 0.600 |
|  | *P* value | 0.014 |
| *DUSP4* | Spearman's correlation coefficient | 0.588 |
|  | *P* value | 0.017 |
| *EPHA2* | Spearman's correlation coefficient | 0.550 |
|  | *P* value | 0.027 |
| *SPRY4* | Spearman's correlation coefficient | 0.144 |
|  | *P* value | 0.594 |

**Figure 2.**
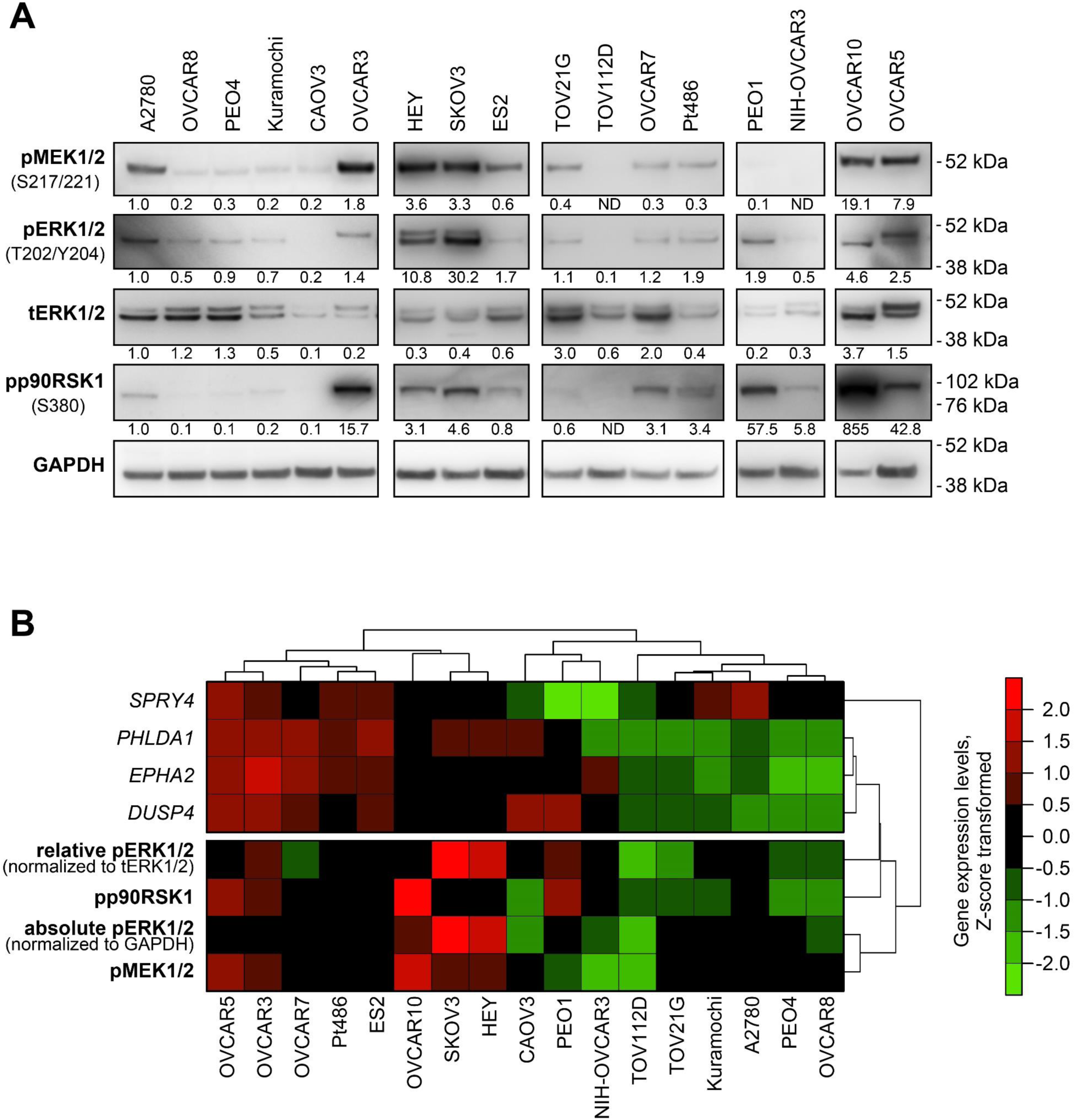
Baseline MEK/ERK pathway activity and *PHLDA1, DUSP4, EPHA2*, and *SPRY4* expression levels in ovarian cancer cell lines. (A) Immunoblotting analysis of phosphorylation of MEK/ERK pathway components in 17 human ovarian cancer cell lines. The image represents data obtained from several separate membranes. An A2780 sample was used in each membrane as a reference sample (see Figure S4). Numbers under the bands represent relative intensity normalized to GAPDH levels and the A2780 sample. (B) Cluster analysis of associations between baseline expression levels of MEK/ERK responder genes and levels of phosphorylated proteins involved in MEK/ERK pathway. Gene expression levels estimated via RT-qPCR and protein levels estimated via immunoblotting were subjected to Z-score transformation. Unsupervised hierarchical clustering was performed using 1-minus Spearman’s rank correlation metrics.

### COMS genes are suitable for monitoring long-term, but not short-term transient changes in MEK/ERK activity

The level of MEK/ERK pathway activity (defined either via direct protein assessment or biomarker signatures) may define the response of cancer cells to MEK inhibition (26, 29). We therefore tested whether the modulation of MEK/ERK activity and COMS genes during cell cycle progression is associated with trametinib effects on cell proliferation. PEO4 cells enriched in different phases of the cell cycle were subsequently treated with trametinib for 72 hours. Cells displaying high pERK1/2 levels (Figure 1C-D) at the moment of trametinib administration responded significantly more robustly to anti-proliferative effects of MEK inhibition (Figure 3A). Phosphorylated ERK1/2 levels exhibited strong negative correlations with the number of viable cells after trametinib treatment, but we observed no significant correlation between expression levels of COMS genes and either pERK1/2 levels or cell number changes (Table S4).

**Figure 3.**
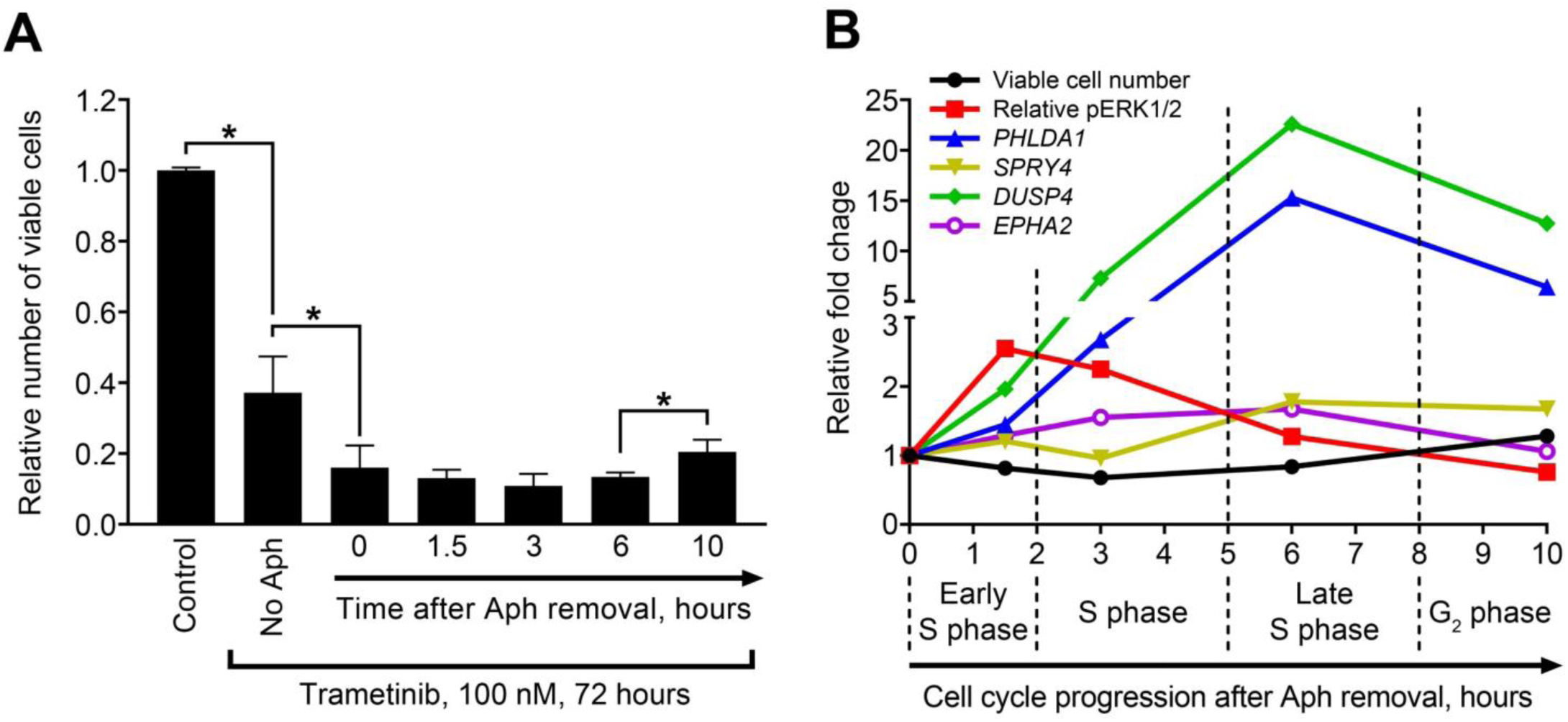
Temporal dynamics of sensitivity to MEK inhibition, ERK1/2 phosphorylation, and MEK/ERK responders’ expression during cell cycle progression. (A) Numbers of viable PEO4 cells enriched in different cell cycle phases after subsequent treatment with trametinib. Data are normalized to control samples and presented as mean±S.D. N=3. * P<0.05, Kruskal-Wallis H test). Aph – aphidicolin. (B) Comparison of trametinib sensitivity, ERK1/2 phosphorylation, and gene expression dynamics across different stages of cell cycle. Data are normalized to 0 hour samples and presented as mean values. Relative pERK1/2 levels were calculated by normalization to total ERK1/2 levels.

To examine more complex connections between these parameters, we compared the kinetic curves of viable cell numbers and pERK1/2, *PHLDA1, DUSP4, EPHA2*, and *SPRY4* levels (Figure 3B). ERK activation and trametinib’s impact on cell proliferation were most prominent within 3 hours after aphidicolin release (early and mid-S phase) and then gradually decreased, while upregulation of COMS genes peaked at 6 hours and *PHLDA1* and *DUSP4* levels remained high even at the 10-hour timepoint. Due to such a postponed response, we concluded that COMS responder genes are more suitable for estimating lasting activation of the MEK/ERK pathway (due to either prolonged external stimulation or long-term internal effects) than for monitoring transient modulations in MEK/ERK activity.

### COMS genes are overexpressed in HGSOC samples with high pERK1/2 levels

To evaluate the potential of COMS genes to predict baseline MEK/ERK activity in clinical samples, we used proteomic and transcriptomic data for large sets of HGSOC cases available from the Nature 2011 and PanCancer Atlas datasets in The Cancer Genome Atlas (TCGA). First, we analyzed 414 samples with RPPA-based proteomics data using an unsupervised hierarchical clustering algorithm and identified three cohorts of patients exhibiting low (n=72), moderate (n=128), and high (n=214) levels of pERK1/2 protein (Figure 4A). Kaplan-Meyer analysis revealed that low pERK and moderate pERK cohorts displayed no significant differences, while the high pERK group had worse overall survival prognosis (Figure 4B). This difference was even more prominent when we merged the low and moderate pERK cohorts into one group (Figure 4C).

**Figure 4.**
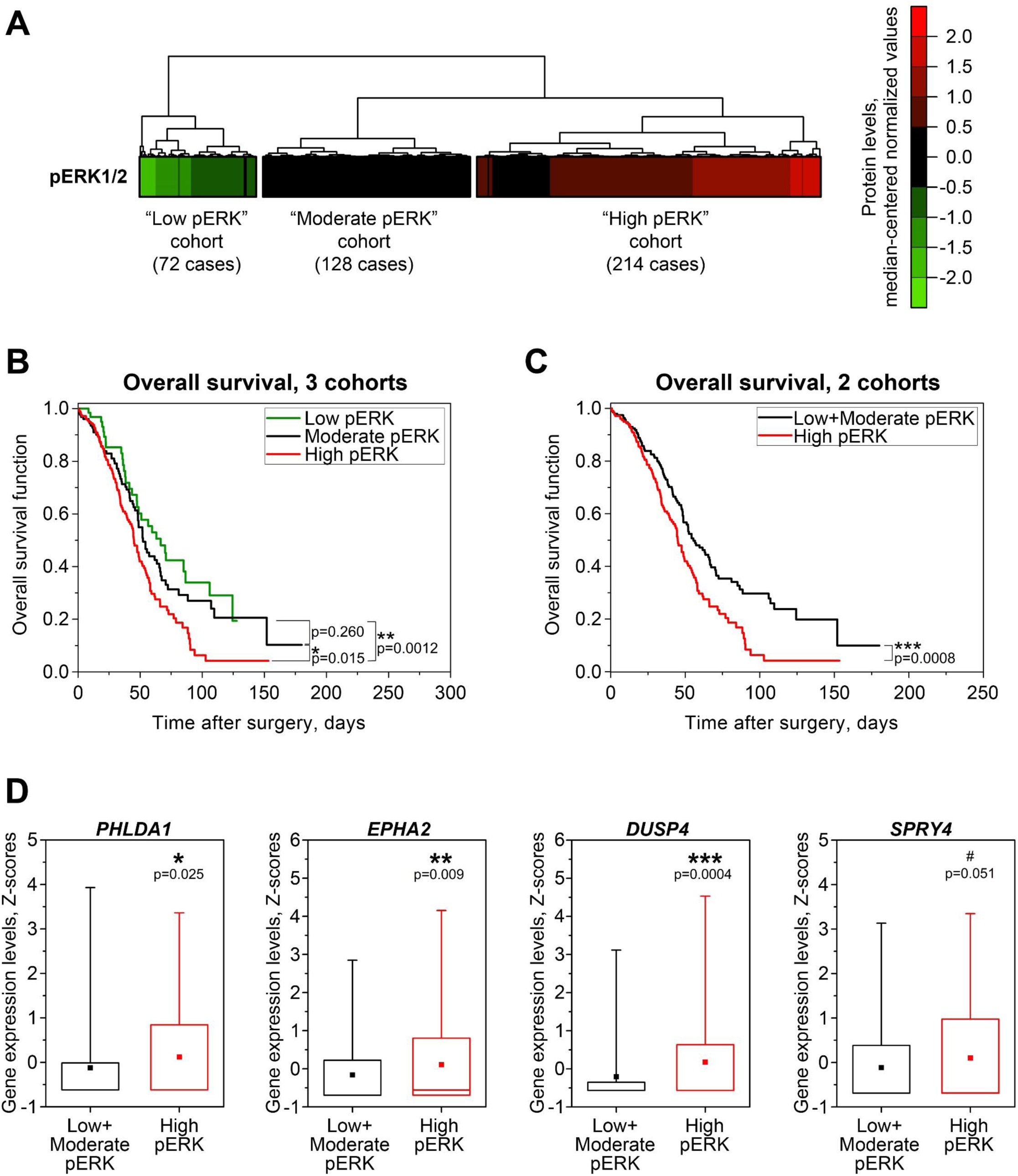
Associations between MEK/ERK pathway activity, patient survival, and MEK/ERK responder genes’ expression in HGSOC patient samples from TCGA. (A) Cluster analysis of phosphorylated ERK1/2 protein levels in samples from TCGA PanCancer Atlas dataset (N=414). Unsupervised hierarchical clustering was performed using Euclidean distance metrics. (B) Kaplan-Meyer analysis of overall patient survival differences between three pERK1/2-defined cohorts from TCGA PanCancer Atlas dataset (N=414). Statistical significance was evaluated using log-rank tests. (C) Kaplan-Meyer analysis of overall patient survival differences between the high pERK cohort and combined low pERK”+”mModerate pERK cohort from the TCGA PanCancer Atlas dataset. Statistical significance was evaluated using log-rank tests. (D) Expression levels of MEK/ERK responder genes in samples from the TCGA Nature 2011 dataset with available pERK1/2 protein data (N=337). Boxes represent median and quartile values, whiskers represent minimum and maximum values, and squares represent sample mean values. Statistical significance was evaluated using two-tailed T-tests with Welch’s correction.

Transcriptomic microarray-based gene expression values were available in the Nature 2011 dataset for 337 cases of 414 PanCancer cases with proteomic data. High pERK samples exhibited statistically higher expression of *PHLDA1, DUSP4*, and *EPHA2* compared to samples from the Low+Moderate pERK cohort. *SPRY4* expression patterns followed a similar trend, but the differences between cohorts did not reach statistical significance, confirming our previous statement about the inferior potential of *SPRY4* as a marker of MEK/ERK response (Figure 4D). We concluded that estimating expression of COMS genes may be used as an additional approach for more reliable evaluation of MEK/ERK pathway functional activity in clinical samples of HGSOC.

### EPHA2 is the only COMS gene with potential to predict patient survival

Since high pERK levels in HGSOC samples were associated with poor overall survival prognosis, and *PHLDA1, DUSP4*, and *EPHA2* were overexpressed in High pERK cases, we evaluated the prognostic potential of these genes using Cox regression models. Advanced patient age at diagnosis, initial platinum resistance, and high *EPHA2* expression were identified as independent predictors of unfavorable outcome, while other COMS genes did not display any predictive power (Table 3). Based on these results, we conclude that baseline *EPHA2* expression level in HGSOC tissue may affect patient survival, but the overall predictive potential of well-established risk factors (advanced age and platinum resistance) is still superior in comparison to COMS signature expression.

**Table 3.** Cox Proportional Hazards analysis of associations between clinical parameters of HGSOC cases, COMS gene expression levels, and overall survival of HGSOC patients from TCGA datasets.

| Clinical parameters | Hazard Ratio | *P* value |
| --- | --- | --- |
| Platinum resistance | 2.6677 | 0.0001 |
| Age at diagnosis | 1.0323 | 0.0047 |
| *EPHA2* expression | 0.6979 | 0.0078 |
| Fraction of genome altered | 0.3313 | 0.1091 |
| Neoplasm histologic grade | 1.3629 | 0.3165 |
| Aneuploidy score | 1.0128 | 0.4758 |
| *PHLDA1* expression | 1.0602 | 0.5981 |
| Tumor residual disease | 1.0521 | 0.6463 |
| *DUSP4* expression | 0.9917 | 0.9474 |

## DISCUSSION

MEK/ERK pathway activation is one of the most common molecular events occurring in cancer cells (including but not limited to ovarian cancer) and is directly connected to malignant cell proliferation and survival (12, 18). While MEK/ERK signaling is often assessed via immunochemical detection of ERK phosphorylation, accumulating evidence suggests that this approach may not always correctly represent MEK/ERK pathway activity (25, 26). Detection of transcripts regulated by pathway of interest is a viable alternative due to high sensitivity and specificity of PCR- and sequencing-based methods. Several transcriptional signatures associated with functional activity of MEK/ERK pathway in various tissues were reported in the literature (17, 26–29), and a 10-gene “MPAS” signature displayed a highly robust MEK inhibition response across multiple tumor types (29). We hypothesized that MPAS signature may be further optimized for ovarian cancer cases and performed detailed evaluation of expression of MPAS genes in several experimental models and sets of clinical samples of ovarian cancer. Only three out of 10 MPAS genes (*PHLDA1, EPHA2*, and *DUSP4*) displayed reproducible associations with baseline levels and changes in MEK/ERK activity across most model systems, with *EPHA2* being the only biomarker with a prognostic potential. Based on these data, we conclude that the MPAS signature is significantly impacted by tissue specificity, model specificity, and temporal and technical details of experiments.

Identification of universal molecular signatures associated with pathological processes is a goal in biomarker research. Prominent tissue heterogeneity affects the functional properties of even the most ubiquitous genes and therefore should be considered when evaluating any biomarker signature. Genes are usually considered tissue-specific if they are only expressed in selected tissues due to unique combinations of genetic, epigenetic, and protein-related factors (34, 35), but such genes are rarely included in universal signatures. On the other hand, factors involved in tissue-specific loss of gene expression, including gene silencers (36), alternative transcript variants (37, 38), and transcriptional repression (39) may drastically impact the performance of such signatures by rendering their important components unusable. We observed such a situation with the *EPHA4* gene, which was expressed below reliable detection levels in all examined ovarian cancer cell lines. According to the original MPAS report, *EPHA4* contributes significantly to MPAS in lung cancer and melanoma, but is virtually without effect in colorectal cancer, confirming its tissue-specific nature (29). We therefore suggest that MPAS or any other signature characterized in non-ovarian samples should be verified based on specific patterns of gene expression in ovarian cancer. This statement is further supported by the idea of assigning tissue-specific weights to commonly used gene sets, to improve statistical power of gene set enrichment analyses (40).

While all our data were obtained from ovarian cancer samples, we utilized several experimental models to investigate different aspects of MEK/ERK pathway activity changes. On one hand, we performed activation or inhibition of MEK/ERK pathway activity with FGF4 or trametinib treatment, respectively – these model systems reproduce prolonged action of external factors that cause long-term shifts in signaling regulation. On the other hand, we evaluated temporal dynamics of MEK/ERK activity during cell cycle progression – these recapitulate more short-term, intricate, and physiologically relevant regulatory changes. Lastly, we investigated stable differences in baseline MEK/ERK activity induced by development of platinum resistance, representing lasting changes that are still present after the initial stimulus (cisplatin treatment) is withdrawn.

We observed significant variability between different experimental models used. *CCND1, ETV4*, and *ETV5* responded prominently to pharmacologic MEK/ERK inhibition in cell cultures, but their changes in trametinib-mediated HGSOC murine xenografts or different cell line models of MEK/ERK activation were much weaker or even completely absent. On the other hand, *DUSP6* displayed moderate response to MEK/ERK activity modulations in all tested experimental models except trametinib-treated cells *in vitro* (Table 1). Such effects, known as model specificity, can be divided into two groups based on unique effects of: (1) the same stimulus observed in different cells/tissues/animals, and (2) similar stimuli observed in the same *in vitro*/*ex vivo*/*in vivo* model system. Both aspects should be considered when analyzing the impact of supposedly similar treatments (e.g. drugs that share a common structure or target the same molecule) performed in different systems (cell lines, organoids, tissue explants, animals). For example, treatment of colorectal cancer cells with thalidomide and its derivatives pomalidomide and lenalidomide results in markedly different transcriptomic changes *in vitro* and *in vivo* (41). Moreover, while the tested agents were expected to act in a similar way, Gene Ontology analysis revealed no common categories affected by all three drugs in either *in vitro* or *in vivo* models, indicating clear differences in their mechanism of action (41). Another recent study compared gene expression response to the proteasome inhibitors ixazomib and bortezomib in human myeloma cell lines and primary patient-derived cells (42). While *in vitro* and *ex vivo* ixazomib treatment models shared more than 200 commonly affected genes, further comparison to effects caused by bortezomib treatment identified only 10 common genes, although both agents selectively target the same 20S proteasome (42, 43). Due to such model specificity, multiple different gene signatures were described for the same drugs or regulatory pathways, resulting in redundancy and decreased reliability (44). In our comparison of five different experimental models of ovarian cancer, only 4 of 10 MPAS genes displayed consistent and uniform response to MEK/ERK activity modulations, and only 3 (*PHLDA1, EPHA2*, and *DUSP4*) were significantly associated with baseline ERK1/2 activity in both cell lines and patient tissue samples, thereby emphasizing the need to validate gene expression signatures for each type of pathology or condition.

The applicability of biomarker signatures for evaluation of both baseline levels of pathway activity and changes during therapy or disease progression greatly improves their usefulness. While many reported signatures are derived from tumor samples obtained prior to treatment, the signatures based on treatment-induced changes in gene expression displayed superior prognostic potential, most likely due to association of such signatures with acquired aggressiveness and resistance (45, 46).

Platinum resistance is an important prognostic factor for HGSOC patients, and its development is tightly associated with MEK/ERK pathway hyperactivation, which also manifests itself in overexpression of COMS genes (18, 47). On the other hand, robust MEK/ERK responders such as *PHLDA1* and *DUSP4* displayed no correlations with survival among HGSOC patients (Table 3). This fact may be explained by the nature of TCGA samples used in our study, since Nature 2011 and PanCancer datasets are almost exclusively comprised of primary HGSOC cases not treated before sample acquisition. We therefore hypothesize that evaluation of MEK/ERK responder genes may be more useful in recurrent ovarian tumors for choosing optimal second-line treatment regimens. Moreover, assessing the dynamic changes of MEK/ERK responders may be theoretically used for monitoring patient response to treatment. Current PCR-based methods are sensitive enough to reliably estimate gene expression levels using extremely low amounts of material, e.g. isolated circulating tumor cells (CTCs), which are detected in 90% of epithelial ovarian cancer patients (48). Evaluation of COMS genes expression in CTCs may provide a non-invasive way of *in vivo* tracking of drug efficiency, disease progression, and chemoresistance development.

Increase in MEK/ERK activity (and expression levels of COMS genes) in cisplatin-resistant cells was associated with increased sensitivity to MEK inhibition (18), which is consistent with increased impact of trametinib upon ERK^high^ cells enriched in S-phase of the cell cycle (Figure 3). Adding MEK/ERK inhibitors to cisplatin (47), ALDH1A inhibitor (18), or Src inhibitor (22) regimens has been suggested as a way to overcome ovarian cancer therapy resistance. Earlier, we reported that trametinib treatment suppresses expression of MEK/ERK-responding genes in ovarian cancer xenografts (18); it is possible that the efficiency of MEK/ERK-targeting therapy in ovarian and other cancers may be monitored via detection of COMS genes (or other optimized transcriptional signatures) in fine needle aspiration biopsy or CTC samples.

Considering the strong association between ERK1/2 phosphorylation modulation and trametinib sensitivity changes observed during cell cycle progression, it may seem counterintuitive that COMS genes display no significant correlations with either parameter (Table S4). However, our experimental model demonstrates that changes in transcript levels are delayed in comparison to protein changes: the peak level of ERK1/2 phosphorylation level is achieved at 1.5 hours after trametinib administration, while gene expression levels reach their maximum values 4.5 hours later (Figure 3B). This delay most likely arises from the inherent difference between protein-to-protein signaling (phosphorylation, cleavage, etc.) and regulation of gene transcription activity. The former is relatively fast as it does not require synthesis of new molecules, while the latter takes more time because of transcription complex assembly, mRNA synthesis, and transcript processing. On the other hand, the magnitude of gene expression response may be much higher than the magnitude of protein changes (*PHLDA1* and *DUSP4* in Figure 3B) due to the accumulation of newly synthesized transcripts over time. Transcript accumulation is strongly connected to lasting response to stimuli (see Table S3) which manifests in detection of significantly elevated transcript levels long after the MEK/ERK pathway activity returns to baseline levels (Figure 1D-E, Figure S3). Our results suggest that the most potent MEK/ERK responders have a lasting response that is useful for detecting long-term changes in pathway activity, especially in terms of MEK/ERK activation. Due to the same reason, the potential of COMS genes for tracking short-term MEK/ERK activity modulations is limited since they cannot demonstrate rapid downregulation upon decreased ERK1/2 activity. Several MEK/ERK-regulated genes termed “immediate early response genes” (*FOS, BTG2, KLF4*) display prominent but transient upregulation within 1-2 hours after pathway activation (49). These genes may be a better choice for monitoring short-term MEK/ERK activity fluctuations, but their expression levels decline over time even with constitutive MEK/ERK pathway activation, rendering their usefulness for possible clinical applications questionable.

## CONCLUSION

Only 3 of 10 MPAS genes (*PHLDA1, DUSP4*, and *EPHA2*) reflect MEK/ERK pathway activity significantly and consistently across multiple experimental models and clinical tissue samples of ovarian cancer. We therefore propose these genes as preferred targets for indirect evaluation of MEK/ERK pathway activity in ovarian cancer. Upregulation of *PHLDA1, DUSP4*, and *EPHA2* together with MEK/ERK activity is associated with acquired platinum resistance, a crucial negative prognostic factor, thus emphasizing that transcriptional signatures should be preferentially developed with consideration of both baseline and treatment-induced gene expression patterns. Altogether, we conclude that any transcriptional signature, regardless of its robustness and flexibility, should be always validated and optimized for tissue type, tumor type, and experimental model of interest due to prominent tissue and model specificity of gene expression in general.

## Supporting information

suppl figure legents

## ACKNOWLEDGEMENTS

The authors express their gratitude to Dr. Rendong Yang (The Hormel Institute, University of Minnesota, Austin, MN, USA) for his advice on TCGA data processing and to Dr. Tim Starr (University of Minnesota Medical School, Minneapolis, MN, USA) for discussing the experimental results. This study was supported in part by a pilot grant awarded to Dr. Chefetz by the Michigan Ovarian Cancer Alliance and the Liz Tilberis grant from Ovarian Cancer Research Alliance.

## DATA AVAILABILITY

Data from “Nature 2011” and “PanCancer Atlas” datasets used in the present study are available from TCGA Research Network (https://www.cancer.gov/tcga) through cBioPortal website (https://www.cbioportal.org) (31, 32).

## FIGURE LEGENDS

**Figure S1.**
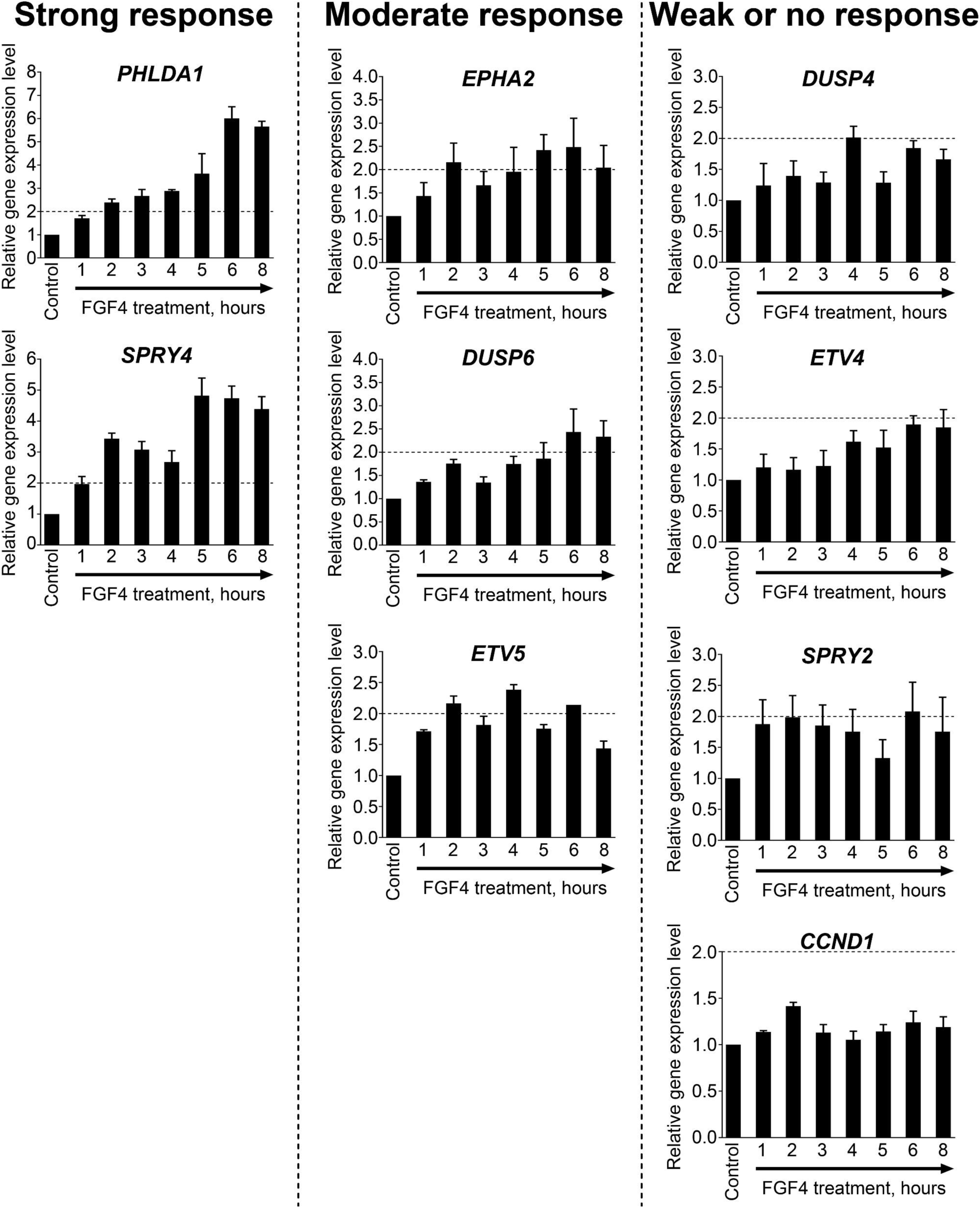

**Figure S2.**
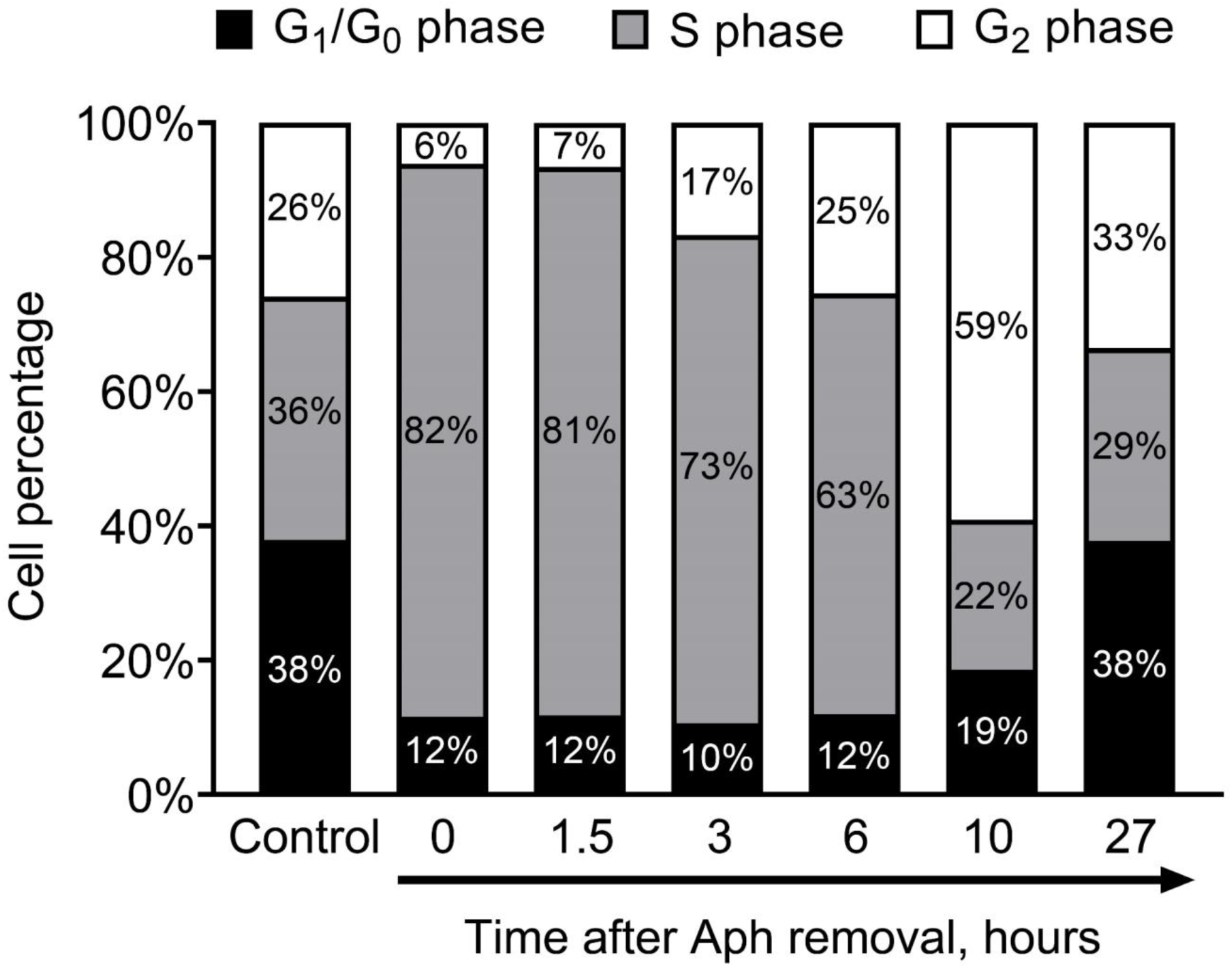

**Figure S3.**
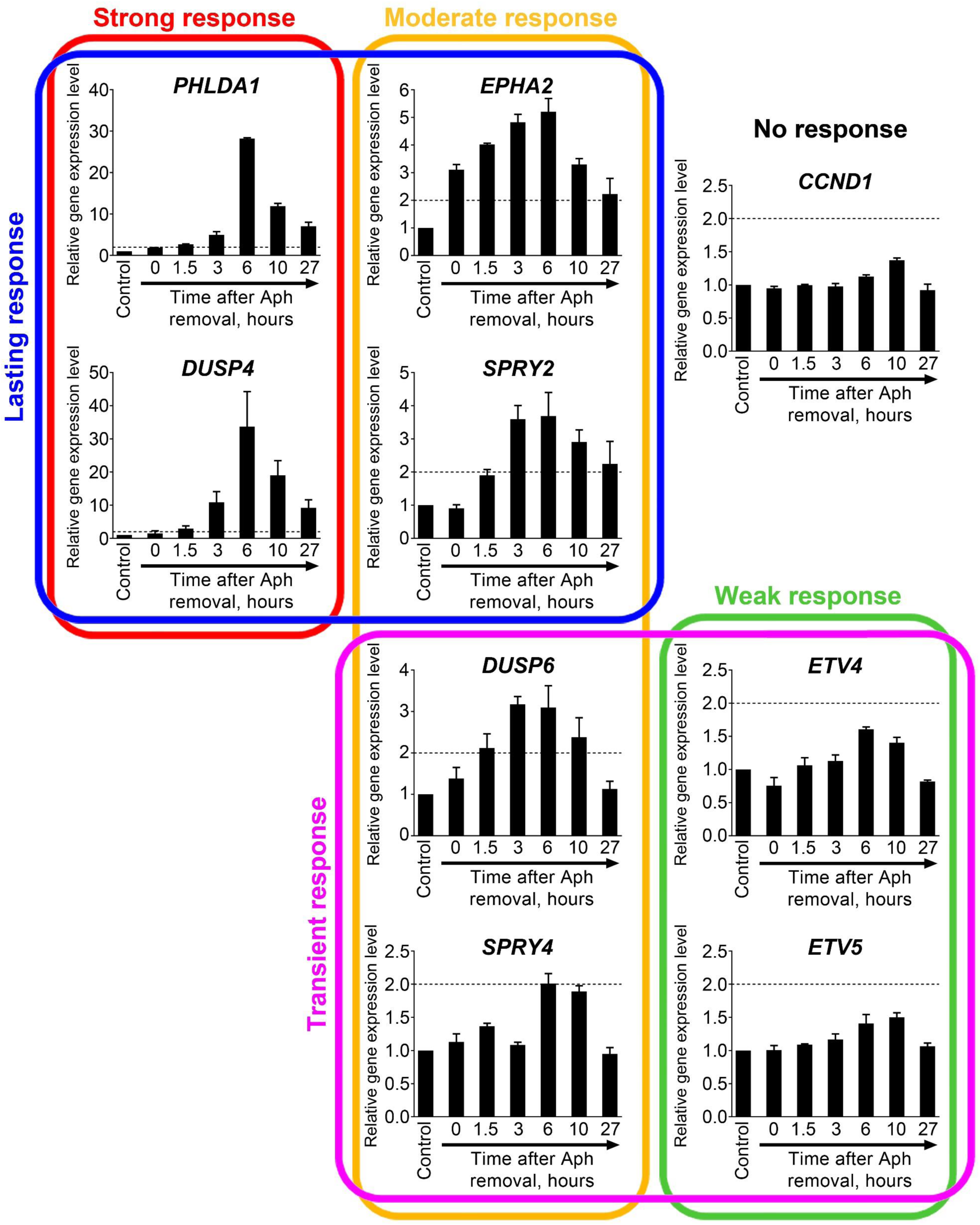

**Table S3.** Summary of MPAS genes’ response to MEK-ERK pathway activation in cell cultures induced by FGF4 treatment or observed during cell cycle progreesion.

| Gene | Response magnitude |  | Response duration<sup>c</sup> |
| --- | --- | --- | --- |
|  | FGF4 treatment<sup>a</sup> | Cell cycle progression<sup>b</sup> |  |
| *PHLDA1* | Strong | Strong | Lasting |
| *EPHA2* | Moderate | Moderate | Lasting |
| *DUSP4* | Weak | Strong | Lasting |
| *DUSP6* | Moderate | Moderate | Transient |
| *SPRY4* | Strong | Moderate | Transient |
| *SPRY2* | Weak | Moderate | Lasting |
| *ETV5* | Moderate | Weak | Transient |
| *ETV4* | Weak | Weak | Transient |
| *CCND1* | No Response | No Response | N/A |

<sup>a</sup>Magnitude of gene expression changes in experiment with FGF4 treatment:
- Strong – more than 4-fold increase; - Moderate – 2-4-fold increase; - Weak – 1.5-2-fold increase; - No Response – less than 1.5-fold increase.
<sup>b</sup>Magnitude of gene expression changes in experiment with cell cycle progression :
- Strong – more than 10-fold increase; - Moderate – 2-10-fold increase; - Weak – 1.5-2-fold increase; - No Response – less than 1.5-fold increase.
<sup>b</sup>Duration of gene expression changes in experiment with cell cycle progression :
- Lasting – gene expression remains upregulated at the end of the experiment; - Transient – gene expression returns to baseline level at the end of the experiment; - N/A – not applicable due to lack of response.

**Figure S4.**
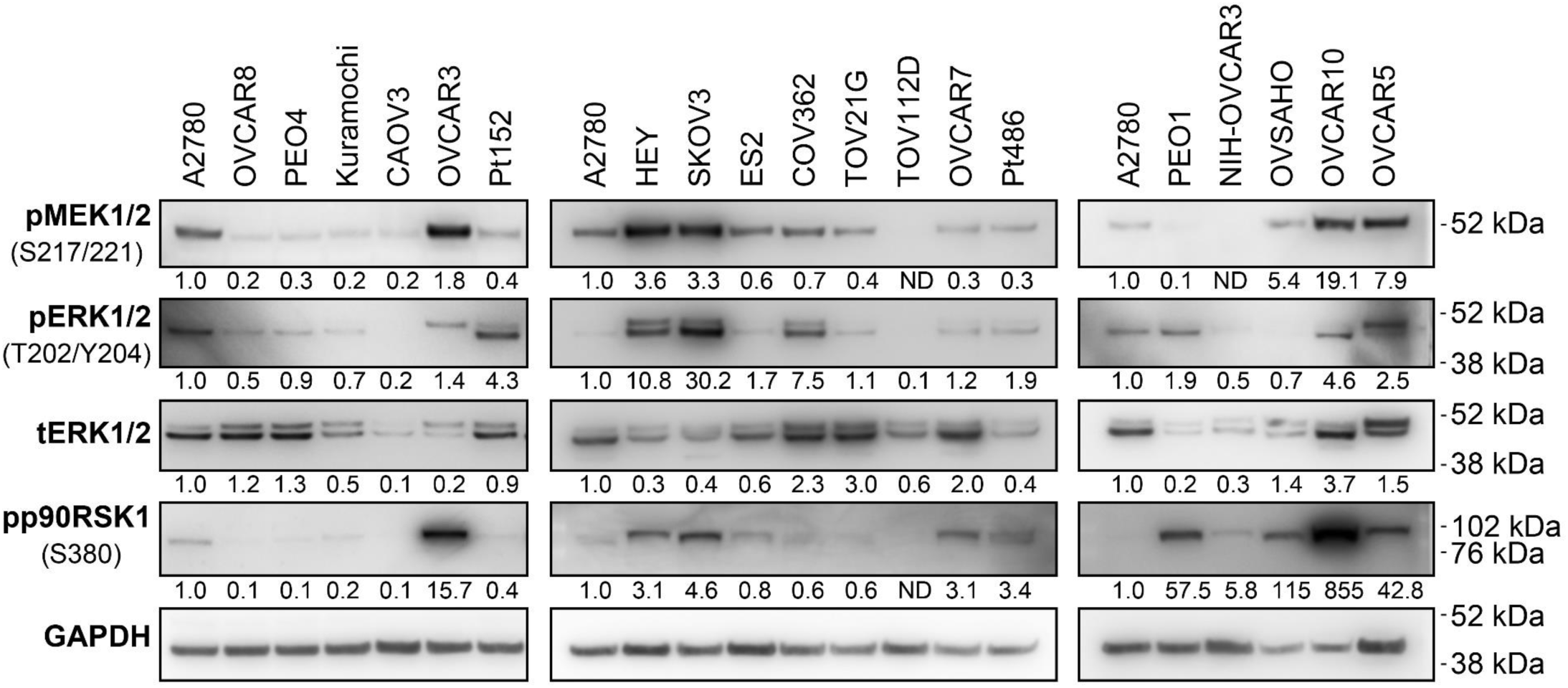

**Figure S5.**
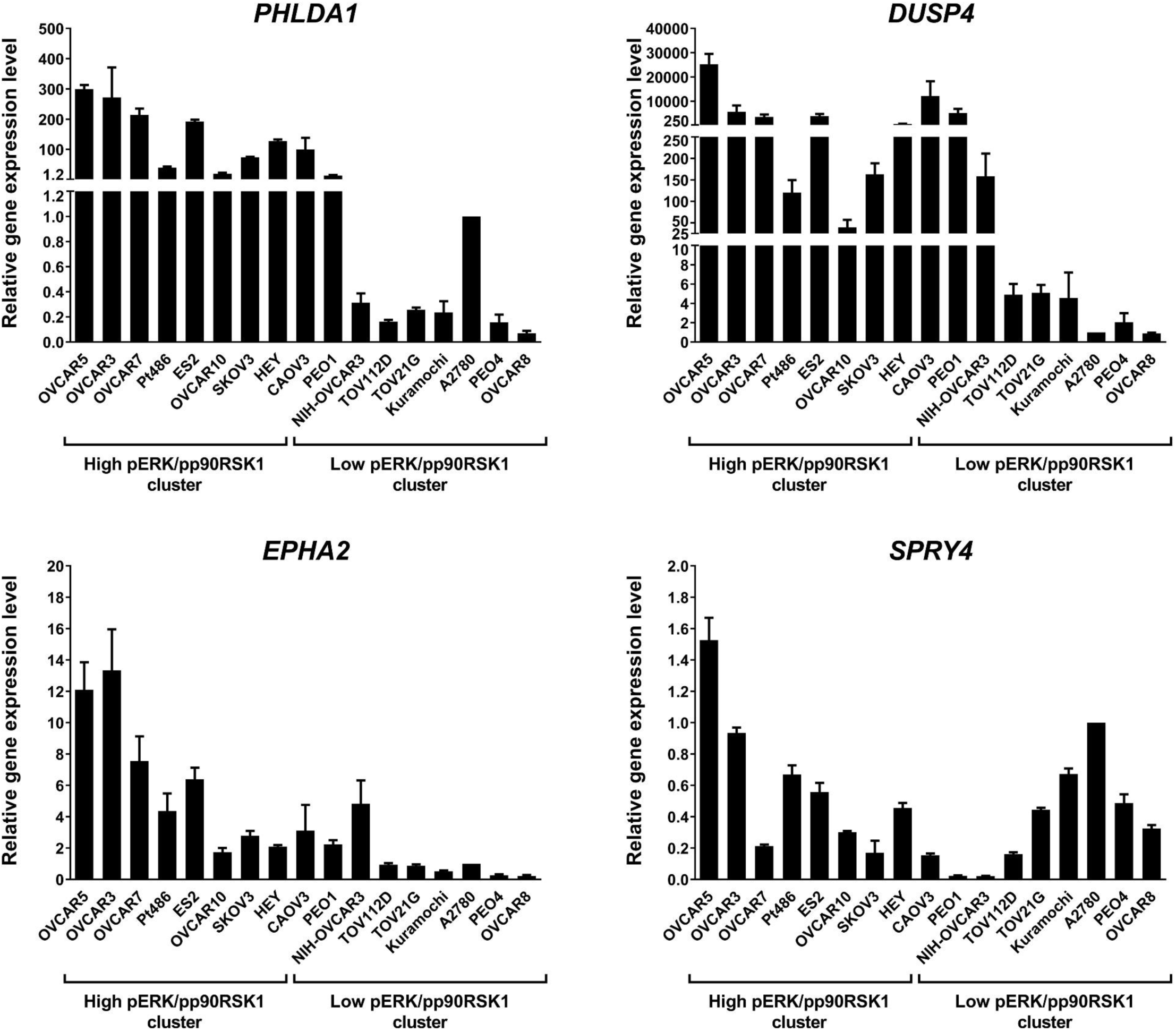

**Table S4.** Spearman’s correlation analysis of associations between ERK phosphorylation changes, COMS genes expression changes, and viable cell numbers in PEO4 cells treated with trametinib during different phases of cell cycle.

| Expression levels of MEK/ERK-responding genes |  | Viable cell number | Relative ERK1/2 phosphorylation level |
| --- | --- | --- | --- |
| Viable cell number | Spearman’s correlation coefficient | 1.000 | **-0.943** |
|  | *P* value | - | **0.005** |
| Relative ERK1/2 phosphorylation level | Spearman’s correlation coefficient | **-0.943** | 1.000 |
|  | *P* value | **0.005** | - |
| *PHLDA1* expression level | Spearman’s correlation coefficient | -0.371 | 0.314 |
|  | *P* value | 0.468 | 0.544 |
| *DUSP4* expression level | Spearman’s correlation coefficient | -0.371 | 0.314 |
|  | *P* value | 0.468 | 0.544 |
| *EPHA2* expression level | Spearman’s correlation coefficient | -0.771 | 0.714 |
|  | *P* value | 0.072 | 0.111 |
| *SPRY4* expression level | Spearman’s correlation coefficient | -0.143 | 0.257 |
|  | *P* value | 0.787 | 0.623 |

