## Supplementary material for "Optimized transcriptional signature for evaluation of MEK/ERK pathway baseline activity and long-term modulations in ovarian cancer": suppl figure legents

**Figure S1. Expression levels of MPAS genes in PEO4 cells treated with FGF4 (100 ng/mL).** Data are normalized to "Control" sample and presented as geometric mean + S.E.M. (N=3, technical replicates).

**Figure S2. The ratios between cell cycle phases in PEO4 cells synchronized with aphidicolin treatment and subsequently released from it.** Aph - aphidicolin.

**Figure S3. Expression levels of MPAS genes in PEO4 cells enriched in different stages of cell cycle.** Data are normalized to "Control" sample and presented as geometric mean + S.E.M. (N=3, technical replicates). Aph – aphidicolin.

**Figure S4. Baseline MEK/ERK pathway activity in ovarian cancer cell lines.** Immunoblotting analysis of MEK/ERK pathway components phosphorylation in 20 human ovarian cancer cell lines. The image represents data obtained from several separate membranes, "A2780" sample was used in each membrane as a reference sample. Numbers under the bands represent relative intensity normalized to GAPDH levels and corresponding "A2780" sample.

**Figure S5. Baseline expression levels of the most robust MEK/ERK responder genes (*PHLDA1*, *DUSP4*, *EPHA2*, and *SPRY4*) in human ovarian cancer cell lines.** Data are normalized to "A2780" sample and presented as geometric mean + S.E.M. (N=3, technical replicates). "High pERK/pp90RSK1 cluster" and "High pERK/pp90RSK1 cluster" are determined based on analysis presented in Figure 2B.
